# Differential neutralization and inhibition of SARS-CoV-2 variants by antibodies elicited by COVID-19 mRNA vaccines

**DOI:** 10.1101/2021.11.24.469906

**Authors:** Li Wang, Markus H. Kainulainen, Nannan Jiang, Han Di, Gaston Bonenfant, Lisa Mills, Michael Currier, Punya Shrivastava-Ranjan, Brenda M. Calderon, Mili Sheth, Brian R. Mann, Jaber Hossain, Xudong Lin, Sandra Lester, Elizabeth Pusch, Joyce Jones, Dan Cui, Payel Chatterjee, Harley M. Jenks, Esther Morantz, Gloria Larson, Masato Hatta, Jennifer Harcourt, Azaibi Tamin, Yan Li, Ying Tao, Kun Zhao, Kristine Lacek, Ashely Burroughs, Terianne Wong, Suxiang Tong, John R. Barnes, Mark W. Tenforde, Wesley H. Self, Nathan I. Shapiro, Matthew C. Exline, D. Clark Files, Kevin W. Gibbs, David N. Hager, Manish Patel, Alison S. Laufer Halpin, Laura K. McMullan, Justin S. Lee, Xuping Xie, Pei-Yong Shi, C. Todd Davis, Christina F. Spiropoulou, Natalie J. Thornburg, M. Steven Oberste, Vivien Dugan, SSEV Bioinformatics Working Group, David E. Wentworth, Bin Zhou

## Abstract

The evolution of severe acute respiratory syndrome coronavirus 2 (SARS-CoV-2) has resulted in the emergence of many new variant lineages that have exacerbated the COVID-19 pandemic. Some of those variants were designated as variants of concern/interest (VOC/VOI) by national or international authorities based on many factors including their potential impact on vaccines. To ascertain and rank the risk of VOCs and VOIs, we analyzed their ability to escape from vaccine-induced antibodies. The variants showed differential reductions in neutralization and replication titers by post-vaccination sera. Although the Omicron variant showed the most escape from neutralization, sera collected after a third dose of vaccine (booster sera) retained moderate neutralizing activity against that variant. Therefore, vaccination remains the most effective strategy to combat the COVID-19 pandemic.

## Main Text

SARS-CoV-2 was first detected in China in December 2019; within six months a variant with a D614G substitution in the viral spike protein became the predominant circulating strain globally. While the D614G variant did not evade antibody-mediated neutralization, enhanced replication and transmissibility of the variant were confirmed in multiple animal models by different groups^1–3^. Enhanced transmissibility and a larger infected population likely led to diversification of the D614G variant into many new lineages. In December 2020, the United Kingdom reported increased transmission of a novel variant of concern (VOC) 202012/01^4^, also referred to as the Alpha (or B.1.1.7, Pango nomenclature) variant^5^. The Alpha variant rapidly disseminated and became the predominant circulating strain in many countries, including the United States (US)^6,7^ (**Fig. 1**). Meanwhile, the Beta (i.e., B.1.351) and Gamma (i.e., P.1) variants were first detected in South Africa in May 2020 and in Brazil in November 2020, respectively, where each variant became the predominant lineage in its respective geographic region^8–10^. By August 2021, the Delta variant (B.1.617.2), which was first identified in India^11^, had displaced the Alpha variant and become the predominant variant within the US (**Fig. 1**) and globally. The Omicron variant (B.1.1.529), which was first reported in late November, is rapidly spreading as of December 2021 and on pace to displace the Delta variant in many regions of the world (**Fig. 1**). The World Health Organization (WHO) and national health authorities, such as the US government SARS-CoV-2 Interagency Group (US-SIG), have designated selected SARS-CoV-2 variants as VOCs or VOIs (**Extended Data Table 1**) based on genomic analysis, transmissibility, disease severity, and, most importantly, impact on the performance of therapeutics or vaccines. Continuous monitoring and rapid characterization of VOCs, VOIs, and other new variants are critical to alleviating the devastating impact of the current pandemic.

**Fig. 1.**
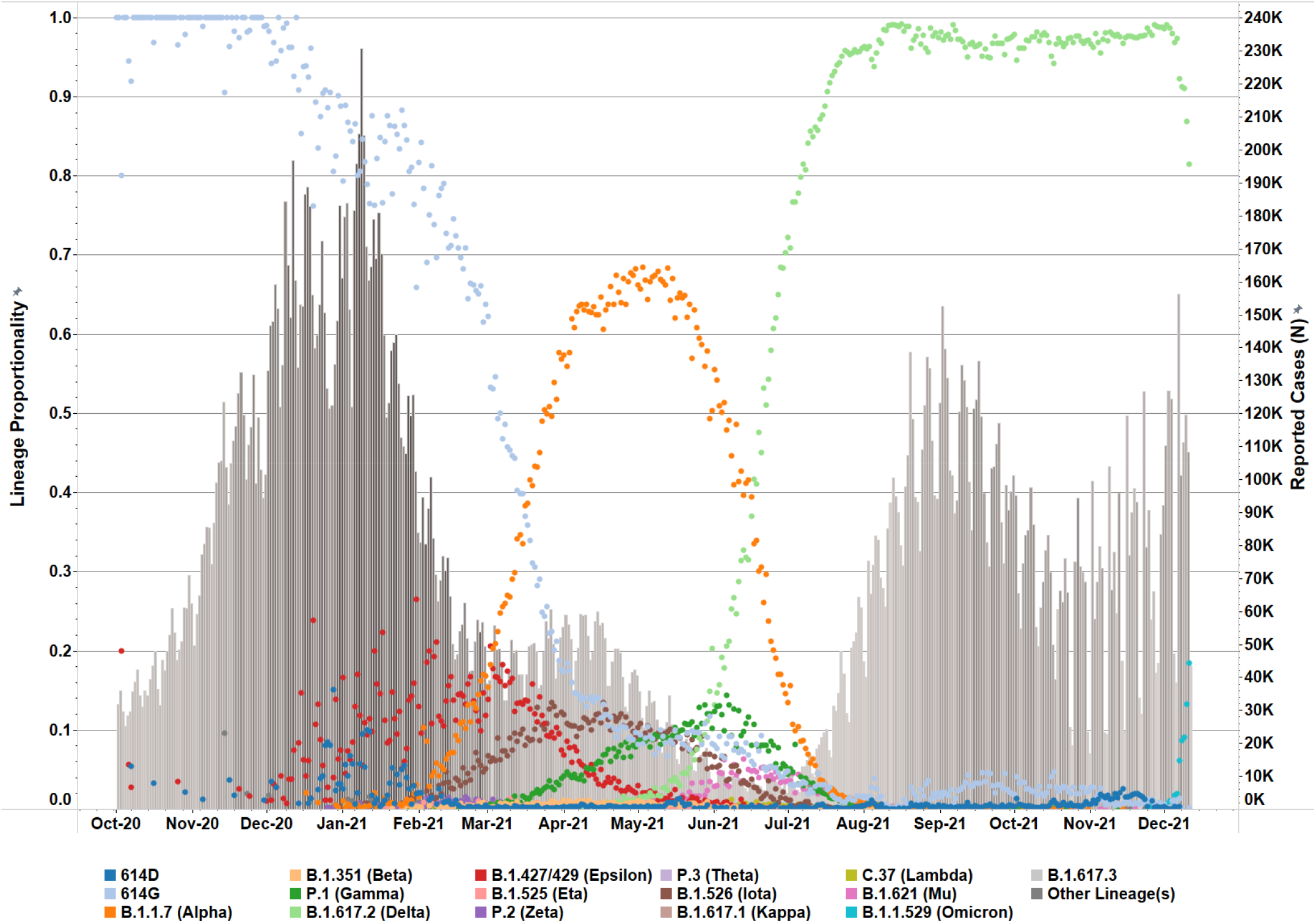
Prevalence of SARS-CoV-2 variants in the United States. SARS-CoV-2 variant prevalence is highlighted for clinical specimens processed within US National SARS-CoV-2 Strain Surveillance networks by relative, daily incidence (dot icons, 0.0 to 1.0) for key Pangolin lineages (WHO nomenclature in parentheses), VOI/VOCs, and specimens encoding critical sequence markers (614D/G) but do not belong to those variants. Daily, reported clinical cases are summarized in the bar graph (right-side, dual Y-axis). Cut-off date is December 12, 2021.

As mRNA vaccines were the earliest and primary form of COVID-19 vaccines administered in the US, we systematically evaluated the neutralization efficiency of U.S. mRNA vaccinee sera against all VOCs and VOIs designated by the WHO Virus Evolution Working Group or US-SIG. To characterize emerging variants in the shortest time frame, particularly in periods which lacked clinical isolates in the US, we generated SARS-CoV-2 fluorescent reporter viruses with VOC and VOI spike substitutions and deletions in the progenitor Wuhan-Hu-1 virus (designated as 614D in this study) by reverse genetics (**Extended Data Table 1**). The reporter SARS-CoV-2 viruses were designed to behave similarly to their clinical isolate counterparts in neutralization assays due to an identical variant spike protein, which is the sole antigen of all vaccines authorized in the US. The spike glycoproteins of many VOI/VOC lineages have subtle differences within the lineages, and the sequence of spike protein used in our studies represent the consensus for the variant lineage or represent a more divergent one from the progenitor within that lineage (**Extended Data Table 1**). For example, the spike of Beta variant tested includes R246I in the N-terminal domain (NTD), which is not found in all Beta lineage viruses.

Using a focus reduction neutralization test (FRNT), we detected minimal impact of the D614G substitution (designated as virus 614G) on the neutralizing activity of the vaccinee sera (fully vaccinated, 2-to-6 weeks post second dose), compared to the progenitor 614D reference virus, of which the spike sequence is most closely related to what was used in mRNA vaccines (**Fig. 2a**). The Alpha variant (B.1.1.7) showed slightly decreased neutralizing antibody titers while the Gamma (P.1), Delta (B.1.617.2 and AY.4.2), Epsilon (B.1.427/B.1.429), Zeta (P.2), Eta (B.1.525), Iota (B.1.526/B.1.526.1), Lambda (C.37), and B.1.617.3 variants showed greater titer reductions but were <4-fold reduced compared to the 614D virus. The Beta (B.1.351), Theta (P.3), Kappa (B.1.617.1), Mu (B.1.621), and Omicron (B.1.1.529) variants showed ≥4-fold reductions in titers, with the Omicron variants showing the greatest escape from neutralization with a 38-fold reduction compared to 614D (**Fig. 2a**).

**Fig. 2.**
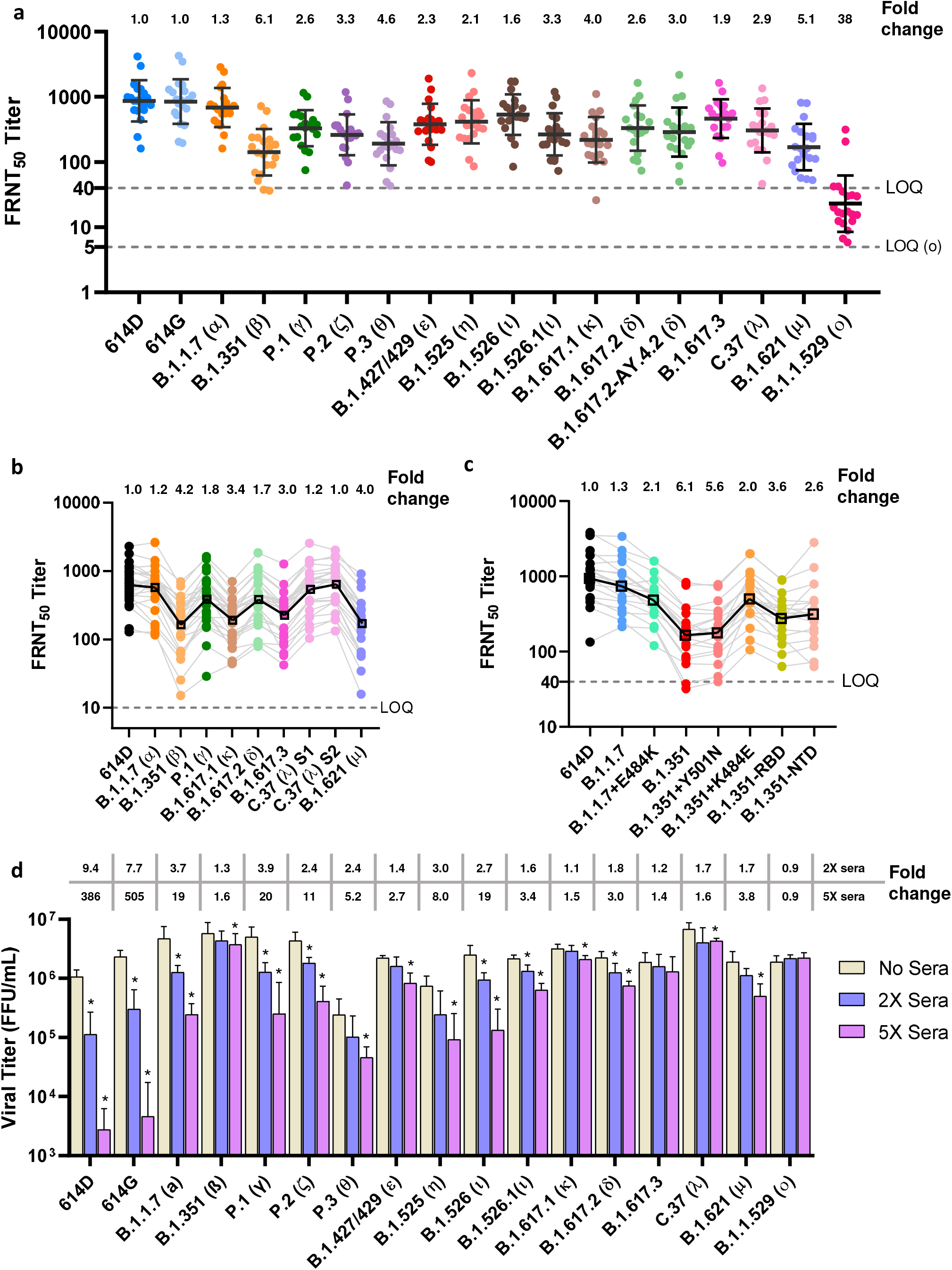
Neutralization and inhibition of mRNA vaccinee sera against live SARS-CoV-2 viruses. Each dot represents the neutralizing titer (FRNT_50_) of an individual serum sample; at least 20 post-second-dose sera were tested against each variant. The average fold changes relative to reference virus 614D (set as 1-fold) are shown on the top of the graph. For each variant, the average fold change is the geometric mean of the individual FRNT_50_ ratios (614D/variant) calculated for each serum sample. Dashed line represents the limit of quantitation (LOQ). **(a)** All WHO and US-SIG designated SARS-CoV-2 variants of concern (VOCs) and variants of interest (VOIs) were tested using reporter viruses. The geometric mean FRNT_50_ titers are shown on the graph with standard deviation. LOD=5 for Omicron and LOD=40 for all other variants. The average fold changes of all variants differ significantly (P<0.0001) from 614D, except for 614G (P=0.9999). **(b)** VOCs and selected VOIs isolated from clinical specimens were tested. The average fold change of all variants differs significantly (P<0.001) from 614D, except for B.1.1.7 (P=0.9995) and the two C.37 viruses (C.37(λ) S1 and S2). **(c)** Reporter viruses with or without specific substitutions were tested to illustrate the impact of specific substitutions. B.1.351 + Y501N has the reversion to original N at 501 of S, and B.1.351 + K484E has a reversion to original E at 484 of S. B.1.351-RBD contains the K417N, E484K, N501Y substitutions in RBD along with the downstream D614G substitution. B.1.351-NTD contains the mutations in NTD along with the downstream D614G. Thin gray lines link the same serum sample tested against the different viruses. The thick black line links the geometric mean titers of different variants. **(d)** Calu-3 cells were infected with 200-400 focus forming unit (FFU) of each virus and incubated for 2 days in media with or without sera. The sera were pooled from the individual sera used in (a) and diluted to 2X or 5X concentration (diluted sera titer FRNT_50_ = 2 or 5 against 614D reference virus). The viruses were collected from Calu-3 supernatant at 2 days post inoculation and titrated by FFU assay. Titer differences are marked as *, representing p<0.05 (One-way ANOVA) for statistical significance, compared to the no sera control within each virus group. Fold changes (reductions) compared to the no sera control are shown on the top of the panel.

As the prevalence of variants rose within the US (**Fig. 1**), the Centers for Disease Control and Prevention (CDC) received an increased number of clinical specimens from state public health laboratories and other CDC collaborating laboratories through the National SARS-CoV-2 Strain Surveillance (NS3) system (https://www.cdc.gov/coronavirus/2019-ncov/variants/cdc-role-surveillance.html). We isolated representative variants from these clinical specimens for characterization by FRNT and sequenced the stocks to ensure the spike correctly represented the appropriate variant lineage. Although the reductions in neutralization differed slightly, they were generally consistent between the reporter viruses and clinical isolates. Prior to the emergence of Omicron, the Beta variant was the most resistant to neutralization, followed by Mu, Kappa and B.1.617.3. The Gamma and Delta variants showed modest escape from neutralization, and the Alpha variant neutralization was not significantly reduced as compared to the 614D reference virus (**Fig. 2b**). Lambda was the only variant that showed a difference between the reporter virus and the clinical isolate, which had an approximate 3-fold reduction versus 1-to-1.2-fold reduction in neutralizing titers, respectively (**Fig. 2a** and **2b**). The Lambda variant has at least 14 substitutions/deletions in the spike (**Extended Data Table 1**) and the limited resistance to neutralization (1-to-1.2-fold) of clinical isolates was surprising. Two Lambda clinical isolates (**Fig. 2b**, Lambda-S1 and -S2) with slightly different spike sequences were analyzed (**Extended Data Table 1**), and the results were consistent. The reporter SARS-CoV-2 system is powerful because the only difference between the viruses being analyzed is the spike, whereas natural isolates have many differences throughout the genome and it’s possible that changes in other gene products (e.g., membrane protein or envelope protein) could impact spike and/or neutralization phenotype but this remains to be understood.

Shortly after the first case of Omicron was confirmed in the US on December 1, we isolated multiple Omicron variants from clinical specimens. Independently, we also collected sera from US vaccinees who received the third dose of a COVID-19 mRNA vaccine (booster) on the day of the booster vaccination (referred as pre-booster sera) and 2-to-6 weeks after the booster vaccination (referred as booster sera). The booster sera had much higher neutralizing titers against the progenitor virus (614D), compared to the post-second-dose sera (**Fig. 2**) or pre-booster sera (**Extended Data Fig. 1**). The booster sera had relatively high neutralizing titers against the Omicron variant at levels comparable to the titers of pre-booster sera against the 614D virus (**Extended Data Fig. 1**). While the post-second-dose sera and pre-booster sera had comparable reductions in neutralization activities (34-to-38-fold reduction), the booster sera had neutralizing activities against the Omicron variant with an average of 19-fold reduction in neutralization titers (**Fig. 2a** and **Extended Data Fig. 1**). Despite the increased neutralization titers detected in booster sera, the substantial reductions in neutralizing titers against Omicron viruses relative to the 614D vaccine reference virus suggests potential for vaccine escape post booster as immunity wanes.

Currently circulating variants are acquiring additional substitutions/deletions, which may further affect transmission, disease severity, or vaccine effectiveness and require prompt evaluation. Utilizing the short turnaround time of reverse genetics, we generated additional Alpha and Beta reporter viruses to examine the effects of specific substitutions that occurred in nature (**Fig. 2c**). While the 3 deletions and 7 substitutions in the Alpha variant spike protein (**Extended Data Table 1**) had a very small impact (1.3-fold) on neutralizing activity of vaccinee sera, we found the addition of the single E484K substitution in spike reduced the neutralizing titer of the Alpha variant by an additional 1.6-fold (2.1-fold compared to the 614D) (**Fig. 2c**). The importance of spike-E484K is further demonstrated in a Beta reporter virus that encoded the spike-484 reversion (B.1.351+K484E), which was found to be 3-fold more susceptible to neutralization compared to an unmodified Beta variant B.1.351 (2.0-fold vs 6.1-fold) (**Fig. 2c**). Interestingly, the N501Y substitution had little impact on the neutralization of the Beta variants (5.6-fold vs. 6.1-fold). A reporter virus containing substitutions only in the Beta variant receptor binding domain (RBD) (B.1.351-RBD, K417N+E484K+N501Y+D614G) was 1.7-fold more susceptible to neutralization compared to the Beta variant with full substitutions/deletions (i.e., 3.6-fold for B.1.351-RBD vs. 6.1-fold for wild type Beta). This result indicates the N-terminal domain (NTD) mutations also contribute to virus neutralization escape. This was confirmed by a Beta variant that only contains mutations in the NTD (B.1.351-NTD, L18F+D80A+D215G+L241-+L242-+A243-+R246I+614G), which reduced the neutralizing titers by 2.6-fold, compared to the 614D virus (**Fig. 2c**). Considering the plasticity of the SARS-CoV-2 spike to substitutions and deletions, as well as the recombination-prone nature of coronaviruses, substitutions and deletions present in current variants may also occur in future variants in different combinations or in different genetic backgrounds. Therefore, it is important to use reverse genetics or other focused approaches to assess the impact and understand the functionality of specific mutations in naturally occurring variants.

To better understand the effect of variant spikes on viral fitness in the presence and absence of neutralizing antibodies, we compared the replication of reporter viruses with the variant spikes in Calu-3 cells, which are a human lung epithelial cell line (**Fig. 2d**). In the absence of inhibitory sera, while Delta replicated to significantly higher titer than the original S-614D reference virus, its infectious titer was comparable to the S-614G virus and many other variants. Interestingly, in the presence of sera, even highly diluted, the titers of 614D and 614G viruses were reduced by more than 7-fold at 2x sera concentration (FRNT50 = 2) and more than 300-fold at 5x sera concentration (FRNT_50_ = 5). As anticipated from neutralization escape data (**Fig. 2a**), the Beta and Omicron variants replicated efficiently in the presence of both concentrations of sera (**Fig. 2d**). Intriguingly, the Delta variant also replicated efficiently and only had a 3-fold reduction in titer at 5x sera concentration (**Fig. 2d**). The Delta variant’s ability to replicate efficiently in the presence of sub-neutralizing concentrations of antisera may have facilitated infections of people with low-to-modest levels of neutralizing antibodies induced by prior infection or vaccination. The immune evasion of Delta variant may be an additive or synergistic result of spike mutations that reduce neutralizing activity and fitness advantages (e.g., better S1/S2 cleavage and cell entry) that are unrelated to neutralization but enhance replication and transmission.

Notably, all post-second-dose sera tested neutralized these variants with FRNT_50_ titers higher than 10 and most of them higher than 40, except against the Omicron variant, which had neutralizing titers between 5 and 40 (**Fig. 2** and **Supplementary Table 1**). It has been widely accepted in the influenza vaccine field that a neutralizing titer (hemagglutination inhibition titer) of 40 or higher is deemed protective (>50% reduction of infection rate)^12^ and a 8-fold reductions in neutralization *in vitro* is a consideration for updating the influenza vaccine strain. For COVID-19 vaccines, few studies on correlates of protection have been published but suggest a titer of 50-100 is the minimum protective neutralizing titer^13,14^ and the fold reduction warranting a vaccine change have yet to be determined. This is also complicated by the unknown role of other effectors in vaccine protection such as memory B-cell or T cell immunity^15^. However, the level of neutralizing antibody titer is apparently predictive of the level of immune protection^16–19^. As neutralizing antibodies wane over time^20^ (**Fig. 2a** and **Extended Data Fig. 1**), infections in fully vaccinated persons by variants circulating at high prevalence are likely to increase. This is especially true for the Omicron variant because the number and location of changes in the S result in considerable antibody escape. Nevertheless, vaccines do prevent and attenuate COVID-19^21^ and anamnestic responses provided through rapid expansion of memory cells should accelerate viral clearance. Therefore, closely monitoring the emergence of variants resistant to neutralization is a necessary and urgent task, and vaccination, including booster doses, remains the most effective strategy to combat the COVID-19 pandemic.

## Acknowledgments

We acknowledge the hundreds of publications each containing neutralization results on some of the VOCs or VOIs reported in this manuscript for which we regret for not being able to cite due to the limitation on the number of references. We thank César G Albariño, Dennis Bagarozzi, Jessica Chen, Jennifer Folster, Matthew Keller, Jimma Liddell, Ji Liu, Gillian McAllister, Magdalena Medrzycki, Krista Queen, Shannon Rogers, Jarad Schiffer, Maria Solano, Sarah Talarico, Brett Whitaker, Jiangwei Yao, and Natosha Zanders for coordination, testing, or analysis support. This work was funded and supported by the CDC COVID-19 Emergency Response and part of the sequencing effort was made possible through support from the CDC Advanced Molecular Detection (AMD) program. Use of trade names is for identification only and does not imply endorsement by the US Centers for Disease Control and Prevention or the US Department of Health and Human Services. The findings and conclusions in this report are those of the authors and do not necessarily represent the official position of the US Centers for Disease Control and Prevention.

## Methods

### Ethics statement

Vaccinee serum samples were collected from individuals through the Influenza and Other Viruses in the Acutely Ill (IVY) Network, a Centers for Disease Control and Prevention (CDC)-funded collaboration to monitor the effectiveness of SARS-CoV-2 vaccines among US adults. Participants had no prior or current diagnosis of infection with SARS-CoV-2 and were fully vaccinated (at least 14 days after the second dose or the third dose) with either Pfizer-BioNTech mRNA vaccine BNT162b2 or Moderna mRNA-1273 vaccine (**Supplementary Table 1**). This activity was approved by each participating institution, either as a research project with written informed consent or as a public health surveillance project without written informed consent. This activity was also reviewed by the CDC and conducted in a manner consistent with applicable federal laws and CDC policies: see e.g., 45 C.F.R. part 46.102(l)(2), 21 C.F.R. part 56; 42 U.S.C. §241(d); 5 U.S.C. §552a; 44 U.S.C. §3501 et seq.

### Biosafety statement

All work involving infectious SARS-CoV-2 virus, including recombinant reporter virus, was performed in CDC Biosafety Level 3 facilities with enhanced practices (BSL-3E). All personnel working with the virus were trained with relevant safety and procedure-specific protocols and their competency for performing the work in the BSL-3E laboratories was certified Recombinant DNA work was approved by CDC’s Institutional Biosafety Committee (IBC). For sequencing, virus was inactivated following protocols approved by CDC’s Laboratory Safety Review Board (LSRB) with a witness confirming that all steps were performed correctly to ensure complete inactivation of virus. After receiving appropriate approvals, inactivated virus was transferred to BSL-2E laboratories for downstream processing.

### Prevalence analysis of variants

SARS-CoV-2 variant statistics for US specimens reported in the National SARS-CoV-2 Strain Surveillance (NS3) and CDC-contracted networks were extracted from a distributed data warehouse and rendered in Tableau Desktop (version 2021.1.1). Daily proportionalities were aggregated by attributed Pangolin (version 3.1.17) lineage assignment including variants of concern (VOC), variants of interest (VOI), and lineages with published World Health Organization (WHO) nomenclature^5^. Pangolin sub-lineages with shared WHO aliases were consolidated: B.1.1.7 and Q sub-lineages (Alpha); B.1.351, B.1.351.2, B.1.351.3 and B.1.351.5 (Beta); P.1 and P.1 sub-lineages (Gamma); B.1.617.2 and AY sub-lineages (Delta); B.1.621 and B.1.621.1 (Mu); and B.1.1.529 and BA sub-lineages (Omicron). Unassigned variants and Pangolin lineages encoding an aspartate (D) or glycine (G) at position 614 were assigned respective “614D” and “614G” labels. Variants that did not satisfy the above criteria were consolidated into “Other Lineage(s).” Clinical statistics included all daily cases and deaths reported to the CDC surveillance network with marked consent. Applied data analytics excluded non-contracted US and global surveillance statistics to limit the impact of non-standardized reporting methodologies and regional over-sampling bias within our dataset.

### Generation of SARS-CoV-2 reporter viruses

#### Risk-benefit analysis

A comprehensive risk-benefit analysis was conducted for using recombinant SARS-CoV-2 reporter viruses in neutralization assays. Briefly, the benefits of using the reporter viruses are: 1) enabling rapid characterization of variants before they are detected in the United States or before CDC receives specimens; 2) eliminating all fixation and staining steps in neutralization assays, shortening the time infectious samples are handled, and reducing chemical safety risks (e.g., formalin) by removing the need to fix cells; 3) minimizing the impact of substitutions in non-spike genes on neutralizing titers, as changes solely reflect the effect of spike mutations; 4) enabling assessment of impact of individual or specific sets of spike mutations; 5) enabling more consistent comparisons as isolates from different clinical specimens were noted to have distinct growth properties even though they were from the same lineage. The associated risk assessments are: 1) reporter viruses are different from any natural virus and created by introducing the spike mutations from a new variant into the backbone virus (progenitor strain Wuhan-Hu-1). The transmissibility of a particular resultant virus could be somewhere between the progenitor virus and the natural variant; 2) as there is limited epidemiological or clinical evidence to suggest spike mutations present in SARS-CoV-2 variants increase pathogenicity, it is most likely the pathogenicity of the reporter viruses will be equivalent or reduced as compared to the progenitor strain or the variant strain; 3) all naturally occurring SARS-CoV-2 variants descending from the progenitor strain have acquired mutations in other genes along with the spike gene. It is possible that some of the non-spike mutations may decrease the transmissibility or pathogenicity of the variant, in which case a reporter virus may be more transmissible or pathogenic than the variant. However, sequence analysis and literature review indicate this risk is very low, especially regarding its potential public health impact during this ongoing pandemic. The safeguard and mitigation strategies are: 1) the backbone of the reporter virus are based on the Wuhan-Hu-1 strain, which is expected to be the least transmissible strain compared to later variants; 2) a mNeonGreen reporter gene replaces the ORF7a in the reporter virus, which may attenuate the virus as the ORF7a protein has been reported to be an interferon antagonist^22,23^; 3) mutations engineered into a reporter virus are either part of or all of the spike mutations found in a natural isolate and the engineering of unnatural mutations is prohibited; 4) the reporter viruses are only to be used in *in vitro* studies, such as neutralization assays, and not in *in vivo* studies; 5) all the *in vitro* work is conducted in BSL-3E facilities including enhanced practices such as shower out after experiments to minimize the possibility of accidental release of the reporter virus to the environment; 6) all staff working with the reporter viruses are fully vaccinated; 7) all staff are approved for working with BSL-3E select agents with senior staff having decades of BSL-3E experience working with highly pathogenic viruses. The **conclusion** is: under the current public health emergency, with the urgency for antigenic surveillance of variants, the benefits of using SARS-CoV-2 reporter viruses exceeds the risks associated with generating and using recombinant reporter viruses. These risks are believed to be extremely low after mitigation.

#### DNA construct

The DNA constructs were either generated as linear fragments as previously described^24^ or generated as a cloned DNA as detailed here. The DNA clone for SARS-CoV-2 strain Wuhan-Hu-1 (GenBank accession number: NC_045512) was purchased from Codex DNA (San Diego, CA). The viral genome was flanked by a T7 promoter sequence at the 5’ end and a linearization site at the 3’ end. The whole cassette was cloned into a bacterial artificial chromosome (BAC) vector. The DNA clone was modified to replace the ORF7a gene with a human codon-optimized mNeonGreen gene (GenBank accession number: AGG56535.1) following the same design as reported previously^24^. The spike gene of this progenitor reporter virus was excised by AscI and BamHI-HF restriction enzymes, resulting in a linearized vector into which synthetic variant spike genes can be assembled using Gibson Assembly (NEB). The Gibson Assembly reaction was then transformed into TransforMax^™^ EPI300^™^ Electrocompetent *E. coli* (Lucigen). Transformations were immediately recovered in SOC medium at 30°C for 1 hour, and plated on LB agar plates containing 25 μg/ml chloramphenicol, followed by approximately 2 days of incubation at 30°C. Colonies were picked and inoculated into LB broth containing 25 μg/ml chloramphenicol for approximately 16±2 hours followed by induction for approximately 4±1 hours at 30°C. DNA was extracted and the sequence was verified by Illumina next-generation sequencing (NGS).

#### *In vitro* transcription

Infectious clones were linearized by SbfI-HF digestion and cleaned up by phenol:chloroform:isoamyl alcohol (PCIA) (25:24:1) extraction. Full-length viral RNA was generated using the T7 RiboMAX^™^ Express Large Scale RNA Production System with slight modifications to manufacturer’s instructions (Promega). Briefly, reaction components were adjusted such that in a 50 μL reaction the final concentration of ATP, CTP, and UTP was 7.5 mM, GTP was 3.5 mM, and the Anti-Reverse Cap Analog (NEB) was used at 2.8 mM. After 2-3 hours of incubation at 30°C, RNA was cleaned up by PCIA and ethanol precipitated for at least 1 hour. Quality of the RNA was assessed by UV-vis spectroscopy and denaturing agarose gel electrophoresis.

#### Nucleocapsid protein expressing cell line

Vero E6 cells (ATCC, CRL-1586) were transfected using Lipofectamine 3000 (Invitrogen) with a plasmid encoding SARS-CoV-2 nucleocapsid protein via CMV3 promoter as well as mCherry2 via an IRES element. Transfected cells were placed under drug selection (0.1-0.3 mg/ml geneticin) to establish the pooled bulk population. Stable single-cell clones were selected from the bulk population by serial dilution plating and drug selection. The expression of nucleocapsid protein was confirmed by the SARS-CoV-2 Nucleocapsid Protein ELISA Kit (ABclonal, Woburn, MA) and the cell clone supporting the most efficient virus rescue was selected (VeroE6-N). Cells were maintained in DMEM supplemented with 10% FBS and 0.2 mg/ml geneticin.

#### Virus rescue

To rescue the SARS-CoV-2 reporter virus, VeroE6-N cells were trypsinized, washed with Opti-MEM (ThermoFisher) and resuspended in 100 μL nucleofector solution at a concentration of 1.5 x 10^6^ cells/100 μl following the instructions of the Nucleofector Kit V (Lonza). *In vitro* transcribed RNA (5 μg) was added to the cells and the cell-RNA mixture was transferred into an electroporation cuvette. Electroporation was completed using the Program T-024 of the Nucleofector 2b device (Lonza). Electroporated cells were immediately transferred into a 6-well plate pre-filled with 2 ml/well of pre-warmed Opti-MEM. At 18-24 hours post-transfection, supernatant was collected (P0) and inoculated onto a monolayer of VeroE6/TMPRSS2 cells^25^ (JCRB1819, JCRB Cell Bank). Twenty-four hours post-inoculation, supernatant was collected to make the seed stock (P1). P1 was propagated in T-150 flasks of VeroE6/TMPRSS2 cells at a multiplicity of infection (MOI) of 0.02-0.1 for 24 hours to make the P2 working stock. The working stock was sequenced as described below.

#### Sequence confirmation

All the SARS-CoV-2 reporter viruses were sequenced by NGS to confirm the sequence of the spike gene. Total RNA was extracted from the working stock of each reporter virus and treated with DNase using the DNase Max kit (Qiagen) following manufacturer’s instructions. Five microliters of resulting clean RNA was used for first- and second-strand cDNA synthesis and library preparation using NEB Ultra II Directional RNA library prep kit for Illumina (New England Biolabs, Ipswich, MA, USA). Libraries were barcoded with unique dual indices synthesized in the CDC Biotechnology Core Facility Oligonucleotide Synthesis Laboratory. Resulting libraries were analyzed for size using the Agilent Fragment Analyzer (Agilent Technologies, Inc., Santa Clara, CA) and quantified using the Qubit 4 Fluorometer (Thermo Fischer Scientific, Waltham, MA). Libraries were normalized to equimolar concentrations, pooled, and sequenced on Illumina NovaSeq 6000 (Illumina, San Diego, CA, USA) using the NovaSeq v1.5 SP Reagent Kit (300 cycles). Demultiplexed reads were processed and assembled using the Iterative Refinement Meta-Assembler (IRMA) on a custom CoV-recombinant configuration^26^. The 614D reporter virus (Wuhan-Hu-1 strain with the ORF7a gene replaced by mNeonGreen) was used as the reference. Reads were filtered for a minimum median phred score (Q score) of 27 and a minimum read length of 80 bases. A Striped Smith-Waterman algorithm was selected for read alignment, and final assembly was performed against the reference sequence matched during read gathering. Amended consensus genomes were created from plurality assemblies by ambiguation of bases with coverage < 20x to ‘N’, and positions with a minor allele frequency (MAF) > 0.2 were given ambiguous nucleotide codes according to IUPAC conventions. Quality metrics were calculated using a count of non-ambiguated amended consensus bases to show proportion of recombinant genome assembled, and average coverage depth across the genome was noted. The full genome sequences of all the viruses are being deposited in GenBank and accession numbers will be provided.

### MSD binding assays

Serum samples were analyzed at 1:100 and 1:5000 dilutions for IgG, IgM, and IgA to SARS-CoV-2 nucleocapsid (N), SARS-CoV-2 S1 receptor binding domain (RBD), and SARS-CoV-2 spike (S) protein (V-PLEX SARS-CoV-2 Panel 2 Kit, Meso Scale Discovery, Rockville, MD), as described previously^27^. Serum antibody levels were calculated using Reference Standard 1 and converted to WHO International Binding Antibody Units (BAU/mL) per manufacturer kit instructions.

### Focus Reduction Neutralization Test (FRNT)

#### Reporter virus-based assay

Serum specimens were heat-inactivated at 56°C for 30 minutes, aliquoted, and stored at −80°C. Each serum sample was serially diluted in 3-fold steps (1:40–1:29,160) in sextuplicate in 96-well round bottom plates. SARS-CoV-2 reporter virus was diluted to 3,200-4,000 focus forming units (FFUs) per ml. Diluted serum samples were mixed with an equal volume of diluted virus and incubated for 1 hour at room temperature (21±2°C). Media from confluent monolayer VeroE6/TMPRSS2 in 96-well tissue culture plates was removed, and 50 μl of the serum–virus mixture was inoculated into each well of cells and incubated at 37°C in a 5% CO_2_ atmosphere for 2 hours. The wells were overlaid with 100 μl of 0.75% methylcellulose in DMEM (Gibco), supplemented with 2% HI-FBS and 1x Pen-Strep and incubated at 33°C in a 5% CO_2_ incubator for 16-18 hours. Plates were scanned using a CellInsight CX5 High-Content Screening Platform (Thermo Scientific) running an ‘Acquisition Only’ protocol within Cellomics Scan Version 6.6.0 (Thermo Scientific, Build 8153). All plates were imaged under equal exposure conditions per channel and under 4x magnification.

Foci were identified and quantified using appropriate ‘Spot Detection’ protocol within Cellomics Scan Version 6.6.2 (Thermo Scientific, Build 8533). Spot counts for each channel were exported for further analysis in R (Version 4.0.3). FRNT_50_ values were calculated by fitting the three-parameter log-logistic function (LL.3) to the FFU counts paired with corresponding dilution information. In cases where the Hill Constant was fit at less than 0.5, e.g., incomplete neutralization, FRNT_50_ values were estimated with a two-parameter fit while fixing the Hill Constant to 1. The R script has been deposited in GitHub: https://github.com/CDCgov/SARS-CoV-2_FRNTcalculations/.

#### Clinical isolate-based assay

SARS-CoV-2 isolates were propagated on Vero/TMPRSS2 cells. All stocks were inoculated at multiplicity of infection (MOI) of approximately 0.004 or 0.01 (for Omicron viruses) and harvested at 2 days post-inoculation. The viral spike sequences were verified using unbiased NGS sequencing (KAPA HyperPrep library kit with RiboErase, followed by Illumina sequencing). All cells and virus stocks tested negative for mycoplasma using MycoAlert Plus reagents (Lonza). Heat-inactivated serum samples were serially diluted in 3-fold steps in DMEM supplemented with 2% heat-inactivated fetal bovine serum (HI-FBS), 1x Pen-Strep and sodium pyruvate (Gibco). The serum dilutions were mixed with an equal volume of virus in the same medium (final serum dilutions 1:10-1:7,290) and incubated for 1 hour at 37°C. Vero/TMPRSS2 cells growing in 96-well imaging plates were then inoculated in triplicate with 40 μL of serum-virus mixtures and incubated for 1 hour at 37°C with periodic shaking of the plates. Inocula were removed and cells overlaid with 1.5% medium viscosity carboxymethylcellulose (Sigma-Aldrich) in MEM (Gibco), supplemented with 4% HI-FBS, 1x Penicillin-Streptomycin (pen/strep), and sodium pyruvate. Twenty hours later, the overlay was washed off with PBS, and cells fixed with 10% neutral-buffered formalin, permeabilized with 0.5% Triton X100 in PBS, blocked with 1% bovine serum albumin in PBS, and stained using SARS/SARS-CoV-2 Coronavirus Nucleocapsid Monoclonal Antibody (Invitrogen MA5-29981) as the primary antibody followed by Alexa647-conjugated secondary antibody (Invitrogen). The monolayers were imaged using a BioTek Cytation3 instrument and virus foci (approximately 100-200/well in no-serum control wells) were counted using Gen5 software. The foci counts were normalized to no-serum controls, and 4 parameter nonlinear regression analysis with bottom constraint set to 0, and top value set to 1 (GraphPad Prism v7.04) was used to fit a curve to the data and to determine the FRNT50 value.

#### Data processing and statistical analysis

Geometric mean titers (GMTs) of each virus were calculated using the FRNT_50_ neutralizing titers of all the serum samples tested against that virus. Average fold change of a variant against the 614D virus (Wuhan-Hu-1 or WA-1) was calculated as the arithmetic mean of the corresponding FRNT_50_ ratios (614D/variant) of each serum sample. SARS-CoV-2 variants isolated from clinical specimens were tested upon availability in parallel with WA-1. All serum/variant combinations were tested twice in independent experiments. In Figure 2B, the arithmetic mean WA-1 FRNT_50_ value for each serum is presented (2-8 independent runs). For other variants, the FRNT_50_ fold-differences to WA-1 were determined from two independent runs per serum sample, and the FRNT_50_ value resulting from this fold average and the grand average FRNT_50_ for WA-1 are depicted. For statistical analysis, normality test and residual diagnostics were performed on the data as the assumption of normality was violated, the data were analyzed using a nonparametric test in the SAS NPAR1WAY procedure. The Dwass, Steel, Critchlow-Fligner method^28–30^ was used for multiple comparisons. Statistical analyses were performed using SAS 9.2 (SAS Institute), with p-value < 0.05 considered significant.

#### Virus replication in Calu-3 cells

Calu-3 cells (Human lung epithelial cell line) were obtained from CDC’s Division of Scientific Resources (DSR). Cells were seeded in 12-well plates and cultured 4 to 5 days until cell confluence reached 80-90% before infection. Culture media was removed from the cells before infection and 200-400 focus forming unit (FFU) virus was added into each well (triplicate wells for each virus). The plates were incubated at 37°C in a 5% CO2 atmosphere for 1 hour. Ten individual vaccinee serum samples from persons received Pfizer-BioNTech mRNA vaccine BNT162b2 and 10 vaccinee serum samples from persons received Moderna mRNA-1273 vaccine were each normalized to 500 FRNT_50_ (against 614D reference virus) and pooled separately (500X stock). Pooled Moderna or Pfizer sera was diluted to 2X or 5X concentration (FRNT_50_=2 or 5 against 614D reference virus) in infection media (DMEM supplemented with 2% HI-FBS and 1x pen/strep). The inoculum was removed from each well after incubation and 1ml of infection media with or without the diluted sera was added to corresponding wells (0X, 2X, or 5X FRNT_50_) and the plates were returned to the 37°C, 5% CO2 incubator for further incubation. Two days later, the culture supernatant was collected and titrated by FFU assay. The FFU assay was performed similarly to the FRNT assay by serial dilution of the virus and without mixing the virus with any sera. The foci acquisition and quantification steps were same as described in the FRNT assay. For each variant, the viral titers in the presence of sera were compared to those in the absence of sera to calculate the fold of change (reduction) in titers. The significance of the reduction was analyzed by one-way ANOVA with Dunnett’s multiple comparisons test (no sera vs. 2X sera; no sera vs. 5X sera).

---

**Extended Data Fig. 1.**
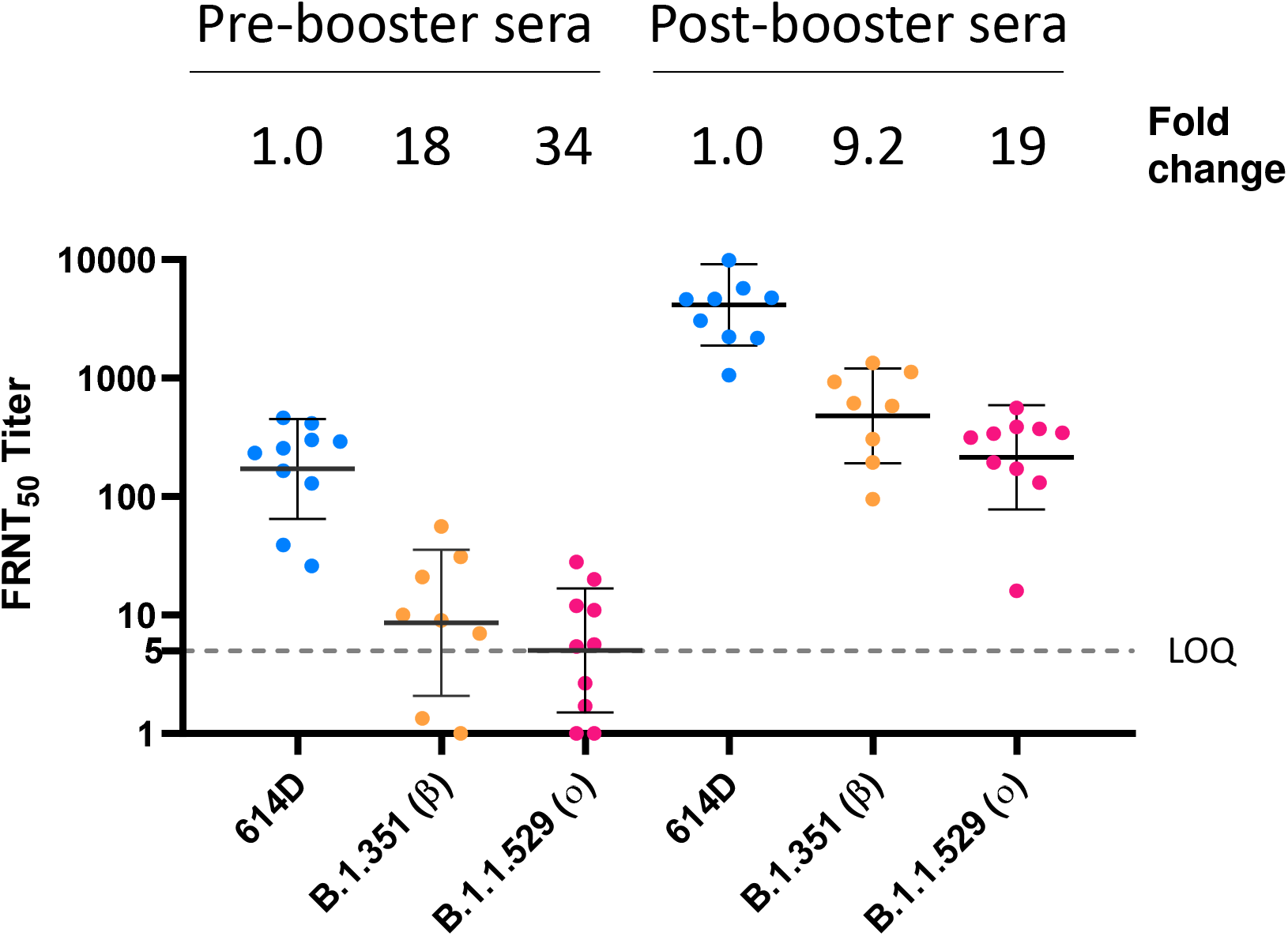
Neutralization activity against live SARS-CoV-2 viruses of mRNA vaccine sera collected pre- and post-booster vaccination. Pre-booster sera were collected on the day of booster vaccination and post-booster sera were collected 2-to-6 weeks post booster vaccination. Viruses were isolated from clinical specimens. The average fold changes relative to reference virus 614D (set as 1-fold for pre-booster and post-booster, respectively) are shown on the top of the graph. For each variant, the average fold change is the geometric mean of the individual FRNT_50_ ratios (614D/variant) calculated for each serum sample. Dashed line represents the limit of quantitation (LOQ). For samples with titer below LOQ the values are still shown and used in calculating the fold change, but they should be interpreted as qualitative instead of quantitative.

**Extended Data Table 1. List of sequence-confirmed substitutions and deletions present in the spike protein of the viruses used in this study.**

**Supplementary Table 1. Quantification post-second-dose sera for nucleocapsid, spike and RBD binding antibody units as well as neutralizing antibody titers against 614D and Delta variant (B.1.617.2).**

